# A genome-wide scan for candidate lethal variants in Thoroughbred horses

**DOI:** 10.1101/2020.05.04.077008

**Authors:** Evelyn T. Todd, Peter C. Thomson, Natasha A. Hamilton, Rachel A. Ang, Gabriella Lindgren, Åsa Viklund, Susanne Eriksson, Sofia Mikko, Eric Strand, Brandon D. Velie

**Author notes:** Correspondence and requests for materials should be addressed to Evelyn T. Todd.

## Abstract

Recessive lethal variants often segregate at low frequencies in animal populations, such that two randomly selected individuals are unlikely to carry the same mutation. However, the likelihood of an individual inheriting two copies of a recessive lethal mutation is dramatically increased by inbreeding events. Such occurrences are particularly common in domestic animal populations, which are often characterised by high rates of inbreeding and low effective population sizes. To date there have been no published investigations into the presence of specific variants at high frequencies in domestic horse populations. This study aimed to identify potential recessive lethal haplotypes in the Thoroughbred horse breed, a closed population that has been selectively bred for racing performance.

In this study, we scanned genotype data from Thoroughbred horses *(n* = 526) for adjacent single nucleotide polymorphisms (SNPs) at high heterozygote frequencies, but with a complete absence of homozygotes. Two SNPs that matched these criteria were mapped to an intronic region in the *LY49B* gene, indicating that a closely linked mutation may cause lethality in homozygous state. Despite a complete absence of homozygotes, almost 35% of Thoroughbreds included in these analyses were heterozygous for both SNPs. A similar loss or absence of homozygotes was observed in genotype data from other domestic horse breeds (*n* = 2030). Variant analysis of whole-genome sequence data (*n* = 90) identified two SNPs in the 3’UTR region of the *LY49B* gene that may result in loss of function. Analysis of transcriptomic data from equine embryonic tissue revealed that *LY49B* is expressed in the trophoblast during placentation stage of development.

In this study, a region in the *LY49B* gene was identified as a strong candidate for harbouring a variant causing lethality in homozygous state. These findings suggest that *LY49B* may have an essential, but as yet unknown function in the implantation stage of equine development. Further investigation of this region may allow for the development of a genetic test to improve fertility rates in horse populations. Identification of other lethal variants could assist in improving natural levels of fertility in horse populations.

**Author Summary:** Recessive lethal mutations may reach high frequencies in livestock populations due to selective breeding practices, resulting in reduced fertility rates. In this study, we characterise recessive lethal mutations at high frequencies in the Thoroughbred horse population, a breed with high rates of inbreeding and low genetic diversity. We identified a haplotype in the *LY49B* gene that shows strong evidence of being homozygous lethal, despite having high frequencies of heterozygotes in Thoroughbreds and other domestic horse breeds. Two 3’UTR variants were identified as most likely to cause loss of function in the *LY49B* gene, resulting in lethality. This finding provides novel insights into the potential importance of *LY49B* in equine development. Additionally, this study may assist with breeding strategies to improve fertility rates in the Thoroughbred and other domestic horse breeds.

## Introduction

There is estimated to be a high rate of natural embryonic mortality in mammals. A large proportion of these embryonic losses occur soon after fertilisation, such that pregnancies often go undetected, with the only sign being reduced fertility (1). Mutation screens in mice reveal that many genes are essential for development, with knockout of 29% of genes tested resulting in embryonic death by day 14 (2, 3). Although mutations in these genes are expected to be under strong negative selection due to being completely deleterious, many species are estimated to carry between one and two recessive lethal mutations per genome (4). However, single mutations are often uncommon in a population, such that unrelated individuals are unlikely to carry the same recessive lethal mutations (5–7).

The likelihood of an individual inheriting two copies of the same lethal mutation is dramatically increased by inbreeding events, whereby alleles that are identical by descent are inherited from a common ancestor (8–10). Populations with high levels of inbreeding and low effective population size (*N_e_*) are at risk of these otherwise rare lethal mutations drifting to high frequencies. Endangered and fragmented species are particularly susceptible due to their small population size and low genetic diversity (11). Recessive lethal mutations can also been found at high frequencies in domestic animal populations despite their large census size due to inbreeding for the selection of desirable traits (12).

In recent years, a number of studies in livestock have identified embryonic lethal mutations at high frequencies due to intensive selective breeding practices (13–20). This is often due to a limited number of sires with desirable characteristics making large genetic contributions to the population (13, 21). Moreover, population bottlenecks due to domestication and breed formation have also resulted in increased deleterious mutation loads and diminished gene pools in many domestic breeds (12, 22–24). These processes lead to genetic variation and *N_e_* being reduced in future generations, which increases the chance of drift and inbreeding events. Lethal mutations that have reached high frequencies are often detected by deviations from the Hardy-Weinberg equilibrium with a lack of homozygotes for one allele (14). Characterisation of such mutations can assist in improving breeding decisions to increase fertility rates in these populations and prevent these mutations from drifting to higher frequencies (25, 26).

The identification of high frequency lethal variants is of particular interest in domestic horse populations. Although a recent study has identified some candidate mutations (27), to date there has been no published comprehensive characterisation of common embryonic lethal alleles in horse populations. Despite the large variety of domestic horse breeds found throughout the world, many breeds suffer from low within-breed diversity and small *N_e_* (28–30). Genetic variation in most horse breeds has decreased markedly within the last 200 years, largely due to their replacement by machinery in agriculture and transport (31, 32). Some horse breeds with large consensus population sizes also experience low *N_e_* and genetic diversity due to intense artificial selective breeding practices and closed population structures (29, 30). Maintaining good fertility rates is particularly important for horse populations due to the seasonal nature of breeding and the low individual fertility output, as mares produce only one foal from an eleven month gestation period (33). Despite the extensive use of hormonal therapies to increase covering success in many domestic horse populations, per cycle pregnancy rates in some breeds only average around 65%, suggesting the presence of unknown variables that may reduce fertility (34).

In this study, we aimed to characterise variants at high frequencies that may cause lethality in the Thoroughbred horse population. The Thoroughbred breed is of particular interest due to the closed population structure since the foundation of the studbook in the 18^th^ century (35). The population has since been intensely selected for the improvement of athletic abilities (36, 37), resulting in contemporary Thoroughbred horses being characterised by high levels of inbreeding and a small *N_e_* (30, 38–40). Due to selective breeding practices, all Thoroughbred horses can trace their ancestry back to a small number of individuals from the foundation of the breed (38, 39). Genetic diversity in the Thoroughbred breed has been reduced in recent decades due to the increased commercialisation of popular stallions providing large genetic contributions to the population (41). Although such practices are in line with selective breeding principles (42), they could also inadvertently increase the frequency of embryonic lethal variants in the population. Reproductive technologies such as artificial insemination are banned in the Thoroughbred population, making the maintenance of high levels of natural fertility imperative. Additionally, Thoroughbred horses have been used as foundation stock for other popular horse breeds including The Quarter Horse, Standardbred, and Warmblood (29), thus identification of embryonic lethal variants in Thoroughbreds is also likely to assist in the breeding management of these populations. Therefore, we also aimed to determine the frequency of any potentially lethal variants identified in the Thoroughbred population in other horse breeds and examine their transcriptomic profile in embryonic tissue.

## Results

### Identifying candidate lethal SNPs at high frequencies in Thoroughbred horses

Analysis of genotype data from Thoroughbred horses (*n* =156) identified only two adjacent, linked SNPs that significantly deviated from the Hardy-Weinberg equilibrium with an absence of homozygotes (Table 1). Under Hardy-Weinberg equilibrium, seven minor allele homozygotes were expected for both of these SNPs in the dataset. Genotype data from Japanese Thoroughbred horses (*n* = 370) also showed an absence of homozygotes for these SNPs (Table 1). In this dataset, the expected number of minor homozygotes for these SNPs under Hardy-Weinberg equilibrium was nine. Despite a complete absence of homozygotes, almost 35% of Thoroughbreds across both of these datasets were heterozygous for this two-SNP haplotype. These SNPs also showed an absence or reduction of minor homozygotes in genotype data for other domestic horse breeds (*n* = 2030, Table 2).

**Table 1:** The allele frequencies of two adjacent SNPs with an absence of minor homozygotes in genotype data from two Thoroughbred horse datasets. The expected number of minor homozygotes in each population was calculated under Hardy-Weinberg equilibrium.

| Population | Sample size | Reference | 6:38278097 |  |  |  | 6:38278874 |  |  |  |
| --- | --- | --- | --- | --- | --- | --- | --- | --- | --- | --- |
|  |  |  | Expected GG | GG | AG | AA | Expected CC | CC | AC | AA |
| Australian Thoroughbreds | 156 | Own data | 7 | 0 | 66 | 90 | 7 | 0 | 66 | 90 |
| Japanese Thoroughbreds | 370 | Fawcett et al., 2019(43) | 9 | 0 | 117 | 253 | 9 | 0 | 117 | 253 |

**Table 2:** Allele frequencies of SNPs in domestic horses at positions 6:38278097 and 6:38278874 on the EquCab2.0 assembly. The total number of horses included in this analysis was 2030.

| Breed | Sample size | Reference | 6:38278097 |  |  |  | 6:38278874 |  |  |  |
| --- | --- | --- | --- | --- | --- | --- | --- | --- | --- | --- |
|  |  |  | Expected GG | GG | AG | AA | Expected CC | CC | AC | AA |
| Swedish Warmblood | 380 | Privately provided, Ablondi et al., 2019(44) | 4 | 0 | 75 | 304 | 4 | 0 | 74 | 306 |
| Norwegian-Swedish Coldblooded Trotter | 646 | Privately provided, Velie et al., 2018(45) | 26 | 0 | 258 | 388 | 28 | 22 | 226 | 393 |
| Quarter Horse | 137 | Petersen et al., 2014(46) | 17 | 0 | 97 | 40 | 17 | 0 | 97 | 40 |
| Exmoor Pony | 285 | Velie et al., 2016(47) | 0 | 0 | 1 | 279 | 0 | 0 | 1 | 282 |
| Various breeds | 582 | Petersen et al., 2013(29) | 15 | 0 | 85 | 497 | 15 | 0 | 85 | 497 |

These two candidate SNPs mapped to the coordinates of 6:38278097 and 6:38278874, which are found in an intronic region of the *LY49B* gene on ECA6. This gene is part of the *LY49* gene family, which plays an important role in innate immunity. There are five functional members of the *LY49* gene family in *Equus caballus,* all of which closely grouped together on chromosome 6. Since both of the SNPs mapped to a non-coding region of the *LY49B* gene, the likelihood of either being a causal variant for lethality is low.

### Phylogenetic origin of the candidate SNPs

According to the phylogenetic tree generated by Petersen et al (28, 29), and their associated SNP data, the SNPs of interest were present in heterozygous state across most phylogenetic branches of domestic horse breeds. Of the 32 breeds in this dataset, 23 had at least one heterozygote for both SNPs of interest. Notably, this two-SNP haplotype was not found in genotype data from one branch of the tree which contains the North Swedish Horse (*n* = 19), Norwegian Fjord Horse (*n* = 21) and Exmoor Pony (*n* = 24) (Table S1). A larger sample of Exmoor Pony data (*n* = 274, Table 2) found only one heterozygote for this haplotype.

### Frequency of the candidate SNPs in other breeds

Analysis of SNP data from other domestic breeds showed that heterozygotes for the SNPs of interest were at a particularly high frequency in the Quarter Horse population (71%, *n* = 137) (Table 2, Table S2). The proportion of heterozygotes was also high in Swedish Warmbloods (*n* = 380) and Norwegian-Swedish Coldblooded Trotters (*n* = 641), being 20% and 40% respectively (Table 2). Smaller datasets also revealed that Belgian Draft (*n* = 19), French Trotter (*n* = 17), Paint (*n* = 15), Morgan (*n* = 19), Mongolian Paulista (*n* = 19) and Tuva (*n* = 15) breeds may also have a high proportion of heterozygotes for this haplotype in their populations (Table S1).

### Identifying candidate causal variants using whole genome sequence data

To further investigate SNP frequencies in this region, variants were called from wholegenome sequence data of 90 domestic horses. The two SNPs identified in the preliminary analysis showed a complete absence of homozygotes for their minor alleles in these individuals (Table S3). Additionally, a number of variants closely linked to these SNPs were identified (Table 3, Figure S1). Annotation of these loci using SIFT(48) identified three variants that may result in changes to protein structure or expression, so these represent the most likely candidates to cause lethality in homozygous state (Figure 1).

**Figure 1:**
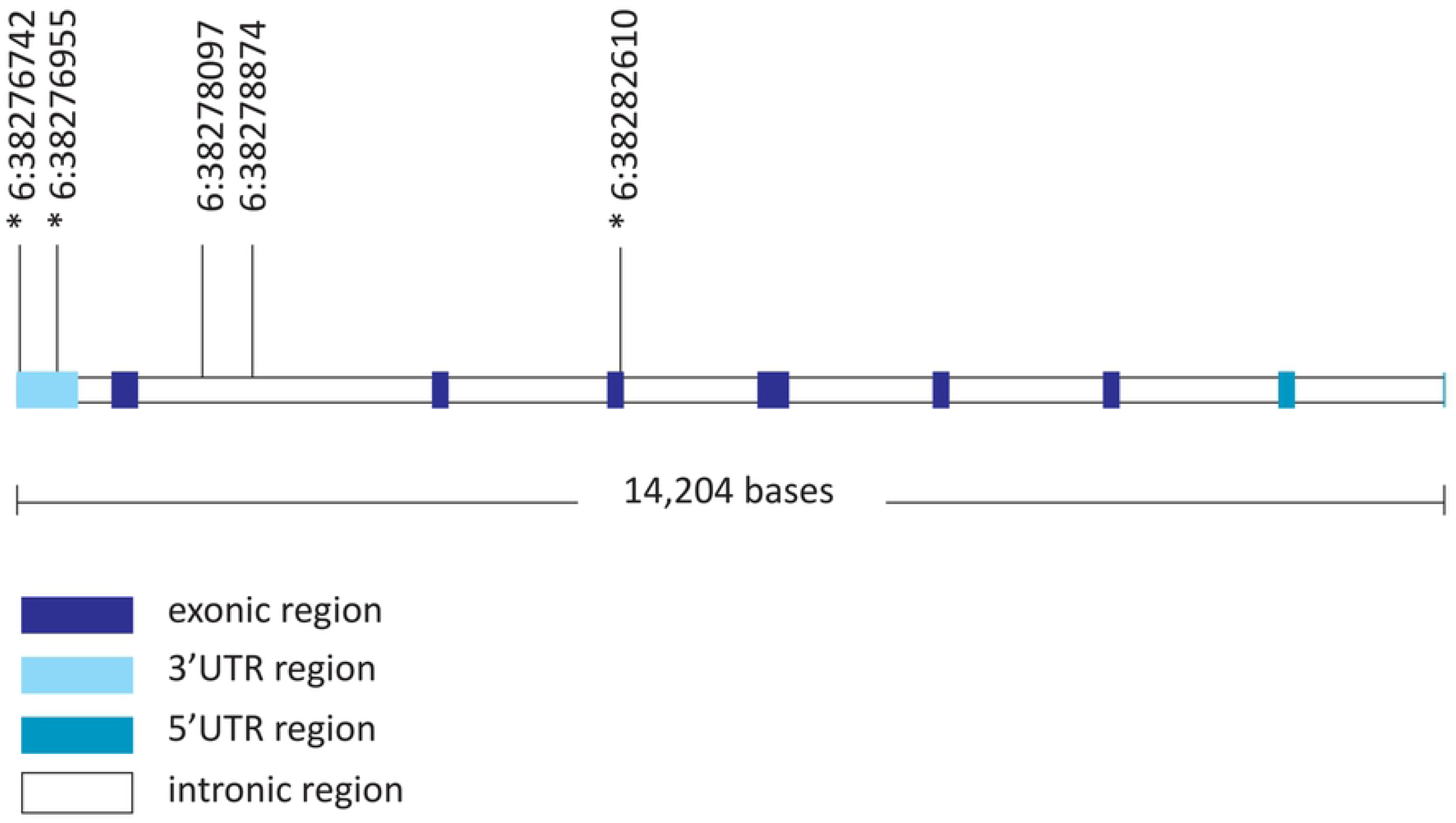
The equine *LY49B* gene structure and SNP positions. The two variants in the intronic region (6:38278097 and 6:38278874) were identified in preliminary analysis as showing a significant absence of homozygotes for one allele. The three variants marked with a * are in linkage disequilibrium to these SNPs and may cause a loss of function in homozygous state. The structure of the gene is based on the EquCab 2.0 reference genome where the *LY49B* gene is on the reverse strand

**Table 3:** Variants linked to the SNPs at 6:38278097 and 6:38278874 on the EquCab2.0 assembly. Variants were identified from whole-genome sequence data for 90 domestic horses of mixed breeds. Only variants within 5 Mb of the SNPs of interest, with a *r*^2^ < 0.8 and a *D*’ < 0.9 were included. Variants most likely to cause a loss of function in homozygous state are indicated with a star.

| Variant | $r^2$ | $D'$ | Distance from<br>6:38278097-6:38278874 | Annotation |
| --- | --- | --- | --- | --- |
| 6:38223012 | 0.83 | 1 | 55085 | Intronic Variant LY49F |
| 6:38264682 | 0.81 | 1 | 13415 | Intronic Variant LY49F |
| 6:38265144 | 0.85 | 0.92 | 12953 | Intronic Variant LY49F |
| 6:38273491 | 0.87 | 1 | 4606 | Intronic Variant LY49F |
| 6:38273498 | 0.87 | 1 | 4599 | Intronic Variant LY49F |
| 6:38273500 | 0.87 | 1 | 4597 | Intronic Variant LY49F |
| 6:38273759 | 0.93 | 1 | 4338 | Intronic Variant LY49F |
| 6:38274367 | 1 | 1 | 3730 | Intronic Variant LY49F |
| 6:38274942 | 1 | 1 | 3155 | Intronic Variant LY49F |
| 6:38276456 | 1 | 1 | 1641 | Intronic Variant LY49F |
| 6:38276742* | 0.83 | 1 | 1355 | 3'UTR variant LY49B |
| 6:38276955* | 1 | 1 | 1142 | 3'UTR variant LY49B |
| 6:38278097 | 1 | 1 | 0 | Intronic Variant LY49B |
| 6:38278874 | 0.93 | 1 | 0 | Intronic Variant LY49B |
| 6:38281733 | 1 | 1 | 2859 | Intronic Variant LY49B |
| 6:38282610* | 1 | 1 | 3736 | Missense variant LY49B |
| 6:38284541 | 0.93 | 1 | 5667 | Synonymous variant LY49B |
| 6:38285848 | 0.85 | 0.92 | 6974 | Intronic Variant LY49B |
| 6:38290614 | 0.85 | 0.92 | 11740 | Intergenic variant |
| 6:38292923 | 0.86 | 0.93 | 14049 | Intergenic variant |
| 6:38294589 | 0.85 | 0.92 | 15715 | Intergenic variant |
| 6:38296564 | 0.85 | 0.92 | 17690 | Intergenic variant |
| 6:38297140 | 0.86 | 0.93 | 18266 | Intronic Variant MAGOHB |
| 6:38297149 | 0.86 | 0.93 | 18275 | Intronic Variant MAGOHB |
| 6:38304295 | 0.83 | 0.91 | 25421 | Intronic Variant MAGOHB |
| 6:38319283 | 0.81 | 0.9 | 40409 | Intergenic variant |
| 6:38334772 | 0.82 | 0.91 | 55898 | Intergenic variant |
| 6:38341477 | 0.86 | 0.93 | 62603 | Intergenic variant |
| 6:38341607 | 0.86 | 0.92 | 62733 | Intergenic variant |
| 6:38345686 | 0.86 | 0.93 | 66812 | Intergenic variant |
| 6:38349116 | 0.86 | 1 | 70242 | Intronic Variant LY49C |
| 6:38349743 | 0.85 | 1 | 70869 | Intronic Variant LY49C |
| 6:38351619 | 0.8 | 0.93 | 72745 | Intronic Variant LY49C |
| 6:38352677 | 0.89 | 1 | 73803 | Intronic Variant LY49C |
| 6:38353265 | 0.85 | 1 | 74391 | Intronic Variant LY49C |
| 6:38354110 | 0.86 | 0.93 | 75236 | Intronic Variant LY49C |
| 6:38358475 | 0.83 | 1 | 79601 | Intergenic variant |
| 6:38360920 | 0.87 | 1 | 82046 | Intergenic variant |
| 6:38399815 | 0.86 | 0.93 | 120941 | Intergenic variant |
| 6:38441535 | 0.8 | 0.93 | 162661 | Intronic Variant LY49E |

The first of these variants, 6:38282610G>A, was located in an exonic region of the *LY48B* gene and resulted in an amino acid change from a phenylalanine to a serine residue. This SNP is located next a tryptophan residue that appears to be highly conserved across members of the *LY49* family and across species. However, there is little conservation of the phenylalanine residue across taxa; some species have a phenylalanine and others a serine at this position. This SNP is annotated as being “tolerated” in SIFT.

Two other variants that were closely linked to the candidate SNPs (6:38276742A>T and 6:38276955G>A) were found within the 3’UTR region of the *LY49B* gene. Alignment of the 3’UTR region of the five functional *LY49* genes in *Equus caballus* revealed that the region containing the SNP 6:38276955G>A is highly conserved in all members of the *LY49* gene family (Table 4). This region may be important for mRNA stability and translation into a functional protein. The other variant was found in an AU-rich region at the end of the *LY49B* mRNA transcript, which is often associated with polyadenylation and post translation stability.

**Table 4:**
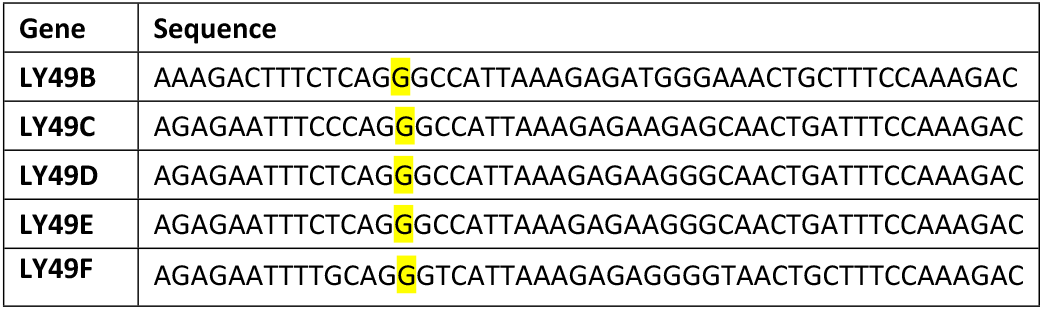
Amino acid residue sequence in a conserved area of the 3’UTR found in all *Equus caballus LY49* genes as mapped in the EquCab2.0 assembly. The SNP position of 6:38276955G>A is highlighted in yellow.

### Transcriptomic analysis of RNA sequence data

Measurable levels of *LY49B* mRNA were not detected in equine trophectoderm tissue collected on day 16 of development. However, *LY49B* mRNA was observed in trophectoderm tissue collected on days 23 and 24 of development (Table 5). Additionally, *LY49B* mRNA transcripts were detected in microarray data from equine chorion and chorionic girdle tissue between days 27 and 34 of development (Table S4). Inner cell mass tissue collected on days 15, 22 and 25 of development did not show any measurable transcription of *LY49B* (Table 5). The genotypes of the candidate SNPs in the mRNA samples analysed were unknown.

**Table 5:** Gene counts from RNA sequence data of three trophectoderm and three inner cell mass tissue samples from equine embryos. Transcript counts are in fragments per kilobase/million (FPKM).

| Tissue | Gene count (FPKM) |  |  |
| --- | --- | --- | --- |
| Trophectoderm | Day 15 | Day 22 | Day 25 |
|  | 0.000 | 0.031 | 0.024 |
| Inner cell mass | Day 16 | Day 23 | Day 24 |
|  | 0.00 | 0.00 | 0.00 |

## Discussion

In this study, we aimed to identify variants at high frequencies in the Thoroughbred horse population that may be lethal in homozygous state. Analyses of genotype data from Thoroughbred horses identified only two adjacent SNPs that fit the strict filtering criteria (Table 1). Genotype data from other horse breeds showed a similar reduction or absence of homozygotes for these SNPs (Table 2). Therefore, this two-SNP haplotype is a strong candidate for harbouring a variant that causes lethality in homozygous state.

The SNPs identified in the preliminary analysis mapped to an intronic region in the *LY49B* gene on ECA6. The *LY49B* gene belongs to the *LY49* (Killer cell lectin-like receptor subfamily A) family of receptors, which consists of five functional members in *Equus caballus* (49). Other species (including humans) have a functionally similar, but structurally different gene family called *KIR* (Killer cell immunoglobin line receptors) (50). The *LY49/KIR* gene family are expressed across various types of immune cells, and mediate their function through bindings to MHC-1 (51). The *LY49B* gene is expressed in myeloid cells where it regulates their activity through an inhibitory effect, possibly to prevent their spontaneous activation (52). Despite the important role that they play in immunity, the function of *LY49* genes in development is currently unknown. In humans, incompatibilities between foetal *KIR* and maternal *MHC* (*HLA*) genotypes are associated with an increased risk of miscarriage and preeclampsia (53–55). Additionally, knockdown of *LY49* in mice showed a high rate of implantation failure (56, 57). These findings indicate that *LY49B* may play an important role in maternal/foetal compatibility and implantation success in horses.

Analysis of transcriptomic data found that *LY49B* was first expressed in equine trophoblast tissue during the placental development stage. The first evidence of *LY49B* expression was found on day 23-24 of development (Table 5), during which the glycoprotein capsule surrounding the embryo is broken down and placental tissue starts to develop (58). Measurable expression of *LY49B* was also found in chorion and chorionic girdle tissues between days 27 and 34 of development (Table S4). During this time, trophoblast cells rapidly proliferate to form the chorionic girdle, which then invades the endometrium to form epithelial cups (59). It is possible that *LY49B* is important for successful implantation of the embryo by mediating the action of MHC-1 which is expressed during this time (60, 61). However, it is also possible that loss of function in *LY49B* may result in post-natal juvenile death, which would explain the lack of homozygotes seen in our data. High rates of juvenile death would be more discernible in the population, so loss of function leading to embryonic death seems more likely. Further investigations into the role of *LY49B* in equine development would confirm whether impaired function causes lethality and the stage of development at which this occurs.

Variant calling in whole-genome sequence data from 90 domestic horses further confirmed an absence of minor homozygotes for the two SNPs of interest. Three variants closely linked to these SNPs were also identified in these data as the most likely candidates to cause loss of function in the *LY49B* gene and result in lethality in homozygous state (Table 3). One SNP was a missense mutation in the coding region of the *LY49B* gene that results in the substitution of a negatively charged serine for an aromatic phenylalanine residue. However, lack of conservation of this SNP in *LY49* genes across taxa makes it seem unlikely to be a causative mutation for embryonic lethality. Two other variants found in the 3’UTR region of the *LY49B* gene were also closely linked to the SNPs identified in the preliminary analysis, and seemed more likely candidates to cause embryonic lethality in homozygous state.

The 3’UTR region of a gene is responsible for transcriptional stability through the binding of miRNAs and RNA binding proteins (62). The addition of the polyadenylation tail to the 3’UTR is also essential to ensure proper processing and translation of the mRNA strand (63). Mutations in the 3’UTR region can lead to degradation of the mRNA, resulting in reduced or inhibited translation even when the gene is transcribed (64). Variation in the 3’UTR region of genes are associated with a number of diseases including Huntington’s and breast cancer in humans (65, 66). Additionally, SNPs in the 3’UTR region are associated with production traits in livestock including milk production in cows, muscularity in sheep and obesity in horses (64, 67, 68).

Despite the importance of the 3’UTR region for the mRNA stability and normal expression of a gene, little is known about how specific polymorphisms can affect post-transcriptional processing. This makes it difficult to identify how the 3’UTR variants identified in this study could affect the translation of *LY49B* mRNA into a functional protein. The 3’UTR variant 6:38276955G>A was identified as a possible candidate for embryonic lethality because it is highly conserved between all members of the *LY49B* family (Table 4) in horses, so may play an important role in mRNA stability. The other 3’UTR variant (6:38276742A>T) is found in an AU-rich region at the end of the transcript, so may be important in the addition of the polyadenylation tail. Further examination of the effects that these variants have on post-transcriptional processing would determine if they impact the normal expression of *LY49B* in horses.

Despite an absence of homozygotes, the two intronic SNPs identified in this study were found at high heterozygote frequencies in the Thoroughbred population. Currently, there is no evidence that variation in the *LY49B* gene is associated with phenotypic advantages in horses. However, it is possible that one of the variants linked to these SNPs confers a heterozygote advantage, which could explain why they have reached such high frequencies in the breed. It is also possible that selective breeding practices favouring a limited number of stallion bloodlines are responsible for this potentially lethal haplotype drifting to high frequencies in the Thoroughbred population. This would be most likely to occur if a stallion that made a large genetic contribution to the population was a carrier. A similar instance has recently been documented in cattle, where a lethal mutation at high frequencies was traced back to a sire with an extensive genetic influence on a population (21).

The presence of this potentially lethal haplotype across many diverse breeds of domestic horses indicates that it may not be the result of a recent mutation present only in the Thoroughbred population. Rather, heterozygotes for this haplotype may have been present in pre-domesticated horses as a rare variant, and have become more frequent in some domestic breeds as the result of population bottlenecks due to breed formations, selective breeding practices and potentially a heterozygote advantage. Domestication and breed formation events have been well documented to result in increased deleterious mutation loads in horses and other domestic species (24, 30, 31, 69). A high proportion of heterozygotes for this haplotype were found in some breeds closely related to the Thoroughbred including the Paint, French Trotter, Morgan and Quarter Horse. Notably, over 70% of Quarter Horse samples included in this study were heterozygous for these SNPs. The Quarter Horse has an open stud book, and higher genetic diversity than the Thoroughbred population (46), making the high frequency of a potentially lethal haplotype at first surprising. The Quarter Horse dataset reportedly did not contain full or half siblings (46), but the collection of samples from one geographical area may not fully reflect the diversity of the worldwide population. An average relatedness analysis of these samples noted the large genetic influence of one particular Thoroughbred stallion (46), which may explain the high frequency of heterozygotes observed in this population. However, the extremely high frequency of heterozygotes in this breed may be due to a heterozygote advantage.

The Belgian Draft, Mangalarga Paulista and Tuva breeds also show a high proportion of heterozygotes, but are more distantly related to the Thoroughbred and to each other. Therefore, the high frequency of heterozygotes in these breeds may be due to independent genetic drift events. Heterozygotes for this haplotype were notably absent from one branch of the tree containing small heavy horses from Northern Europe, which are more distantly related to the Thoroughbred. A larger dataset of Exmoor Pony samples from this phylogenetic branch revealed one heterozygote for this haplotype (Table 2). This could be due to a calling error, but it is also possible that these SNPs exist at very low frequencies in these breeds. The small sample size of the genotype data for many individual breeds in this study means that heterozygote frequencies across all subpopulations found throughout the world may deviate from that reported here. However, these data provide an indication of breeds with high proportions of heterozygotes for this region. Overall, our findings suggest that this region shows evidence harbouring a homozygous lethal variant, yet a high proportion of heterozygotes are found across many domestic horse breeds.

In this study, we identified a haplotype at high heterozygote frequencies in the Thoroughbred horse population that is a strong candidate for harbouring a variant causing lethality in homozygous state. Similar analyses on larger datasets in other livestock populations have identified multiple lethal haplotypes, so it is likely that other such variants are present at high frequencies in the Thoroughbred population but were not captured in this study. Additionally, the use of commercial SNP arrays only allows for the identification of variants with high minor allele frequencies in populations. Analysis of larger sample sizes, and using higher density genotype data could allow for identification of other variants associated with lethality in domestic horses. The identification of this potentially lethal haplotype demonstrates the potential implications of heavily favouring a limited number of bloodlines in selective breeding practices. The association of this haplotype with lethality can be used to assist in breeding decisions to improve mating outcomes in the Thoroughbred population. Although there are a high proportion of heterozygotes in some domestic breeds, limiting the use of stallions that are carriers could reduce the frequency of this haplotype in future generations. Additionally, further investigations into a possible heterozygote advantage could assist in understanding the high heterozygous frequencies across many domestic breeds. Further characterisation of lethal haplotypes in other breeds would also assist in breeding management to increase fertility rates in domestic horse populations.

## Methods

### Initial genotyping

Genotype data from a representative sample of Thoroughbreds were used to identify SNPs with a high proportion of heterozygotes, but an absence of homozygotes for one allele. Genome-wide SNP data were generated for 156 Australian Thoroughbred horses by genotyping samples on either the Illumina 70K Chip (65,102 SNPs) (*n* =102) or the Affymetrix 670K Chip (670,796 SNPs) (*n* = 54). Common genotyped SNPs between the two arrays were scanned for deviations from the Hardy-Weinberg equilibrium with an absence of homozygotes for one allele using PLINK (version 1.9) (70). The *P*-values were adjusted using a false discovery rate correction with the R package “qvalue” (71). Since SNPs with an absence of homozygotes could indicate a calling error, the search was narrowed to only include adjacent SNPs that fit such criteria.

The frequencies of the candidate SNPs were then examined in publicly available genotype data from Japanese Thoroughbreds (*n* = 370) typed on the Affymetrix 670K Chip (43)and these were added the Thoroughbred sample. The SNP frequencies were then characterised from genotype data from Swedish Warmbloods (*n* = 380) (44) and Norwegian-Swedish Coldblooded Trotters (*n* = 670) (47) typed on the Affymetrix 670K Chip. Publicly available data from Exmoor Ponies (*n* = 285, typed on the Affymetrix 670K Chip) (47), Quarter Horses (*n* = 137, typed on the Illumina 70K Chip) (72) and horses of 32 different domestic breeds (*n* = 582, typed on the Illumina 50K Chip) (28, 29) were also included in this preliminary scan for SNP frequencies. In these data, raw intensities were plotted to check for calling errors. If potential calling errors were detected, SNPs were recalled using a mixture model fitted with an expectation-maximization algorithm in R.

### Variant discovery and mapping

Publicly available whole-genome sequence data were used to further examine the frequencies of the candidate SNPs identified in the initial genotype analysis, and to identify linked variants. Paired end whole-genome sequence data from 90 horses of different domestic breeds were used in this analysis (Table S5). The whole genome datasets were downloaded from the European Nucleotide Archives (ENA, https://www.ebi.ac.uk/ena) which included horses of different domestic breeds (PRJEB14779, *n* = 70) and additional Thoroughbred samples (PRJNA168142, *n* = 16 and PRJNA184688, *n* = 4) (Table S5).

The SNP array used in the initial genotyping analysis was developed based on coordinates of the EquCab2.0 reference genome. For consistency, we used the EquCab2.0 assembly as a reference for the whole-genome sequence analysis. The raw reads were mapped to the EquCab2.0 reference genome using BWA-MEM algorithm from Burrows-Wheeler Alignment Tool (version 0.7.17) (73). Duplicate reads were flagged using Samblaster (version 0.1.22) (74), and base recalibration was performed using Genome Analysis Toolkit (GATK) (version 4.0.8.1) (75). Variants (SNPs and INDELs (insertions and deletions)) were called using Haplotype Caller and then filtered using the standard hard filtering recommendations in GATK (76). The individual SNPs were then filtered to only include high quality allele calls with an average filtered depth over 10 and a Phred score over 20.

Variants that were linked to the SNPs identified from the genotype data were produced using the LD function in PLINK (version 1.9) with a window size of 5 Mb (70). Only SNPs with an *r*^2^ value of over 0.8 and a *D*’ value > 0.9 were shortlisted. The effects of each SNP on gene structure and function was characterised using SIFT (version 4G) (48). The conservation of variants across taxa was analysed using the NCBI Conserved Domain Database Search (77).

### Transcriptomic analysis

Publicly available RNA sequence data were used to examine expression levels of the genes of interest in embryonic tissue. The data included equine inner cell mass tissue (collected at day 15, 22 and 25, *n* = 3) and trophectoderm tissue (collected at day 16, 23 and 24, *n* = 3) from the Functional Annotation of ANimal Genomes (FAANG) equine biobank (available from ENA under the project name PRJNA223157) (78). Adaptors were trimmed using bbduk from BBtools (version 37.98) (79). Reads were aligned to the EquCab 2.0 genome using STAR (version 2.7.2b) (80). Counts were generated using featurecounts from subread (version 1.5.1) (81), then quantified in fragments per kilobase/million (FPKM) using the R package “edgeR” (82) with the Equus_caballus_Ensembl_94 file used for annotation. Microarray data for chorion (*n* = 19) and chorionic girdle (*n* = 19) tissue collected from horse embryos between days 27-34 of development (83) were also examined for gene expression levels.

### Ethics statement

DNA samples for Australian Thoroughbred horses were collected under approval from University of Sydney Ethics Committee N00-2009-3-5109. Written informed consent to use the animals in this study was obtained from the owners of the animals. The hair samples from Swedish Warmblood horses were originally collected for parentage testing and stored in the biobank at the Animal Genetics Laboratory, SLU so ethics approval was not applicable. DNA samples of Norwegian-Swedish Coldblooded Trotter were collected under approval from the Ethics Committee for Animal Experiments in Uppsala, Sweden [Number: C 121/14]. All other data was downloaded from publicly available repositories.

## Acknowledgements

The authors acknowledge the technical assistance provided by the Sydney Informatics Hub, a Core Research Facility of the University of Sydney.

## Supporting Information

Additional tables and figures.

